# Highly efficient in *vivo* mutagenesis method based on the endonuclease and exonuclease activity of *Bacillus alcalophilus* RecJ (BaRecJ)

**DOI:** 10.1101/2024.02.16.580766

**Authors:** Jixiang Shang, Yanchao Zhang, Zongjun Xu, Zhongtao Sun, Minggang Zheng, Shouqing Zhang

## Abstract

With the continual advancement of technologies such as microbial cultivation, DNA sequencing, bioinformatics, and genetic engineering, methods for microbial breeding have become increasingly diverse. We identified a RecJ enzyme (BaRecJ) with both endonuclease and exonuclease activities from *B. alcalophilus*. Differing from traditional physical-chemical mutagenesis approaches and the methods based on perturbation factor. this research established a novel mutagenesis method utilizing the endonuclease and exonuclease activities of BaRecJ. Mutagenesis of *E. coli* was conducted using the BaRecJ method, followed by screening for rifampicin-resistant mutants, *rpoB* sequencing results demonstrated a broader, more uniform spectrum of mutations and a higher frequency of substitution mutations with this mutagenesis approach. Furthermore, this mutagenesis method was applied to *S cerevisiae*, resulting in mutants with enhanced tolerance to acetic acid and ethanol, exhibiting improved fermentation performance and flocculation abilities Genomic resequencing analysis summarized genes possibly associated with the tolerance of mutants. Therefore, this approach not only holds immense potential in microbial mutagenesis breeding and adaptive evolution but also, when coupled with genomic resequencing, allows for the rapid identification of genetic loci associated with specific traits.

## INTRODUCTION

Biological evolution is driven by new mutations. In most eukaryotes and prokaryotes, approximately one base mutation occurs every billion replications (Lynch, 2010). Additionally, mutation spectrum are a key component of genetic diversity that fuels evolution. Traditional mutagenesis methods produce a narrow mutation spectrum and are carcinogenic. Ultraviolet radiation can produce a wide range of mutation spectra, but its effectiveness is limited by cellular toxicity. To increase mutation frequency and produce a wide range of mutation spectrum, some global in *vivo* mutagenesis methods based on perturbation factor have been developed (Xu et al., 2018; Wu et al., 2022; Badran and Liu, 2015; Asakura et al., 2011; Luan et al., 2013).

A novel class of RecJ nucleases, distinct from the classic RecJ nucleases and derived from *Bacillus alcalophilus*, *Bacillus cereus*, and *Bacillus halodurans*, exhibit not only the exonuclease activity of RecJ but also endonuclease activity (Ma et al., 2021; Wang et al., 2020; Zheng et al., 2020; Srivastav et al., 2019; Makarova et al., 2012). Based on the combined exo- and endonuclease activities of the RecJ (BaRecJ) nuclease from *B. alkaliphilic*, we established a novel global in *vivo* mutagenesis method, differentiating it from current global in *vivo* mutagenesis mechanisms (Luan et al., 2013; Badran and Liu, 2015).

We applied the BaRecJ in *vivo* mutagenesis method to *E. coli*, achieving the broadest mutation spectrum in *rpoB* (Fernandez et al., 2020; Badran and Liu, 2015). For *S. cerevisiae*, using the BaRecJ mutagenesis approach combined with Adaptive Laboratory Evolution (ALE), mutants with high tolerance to acetic acid and ethanol were obtained. Overall, this system is applicable to both prokaryotic and eukaryotic organisms for mutagenesis, significantly enhancing in *vivo* mutagenic capabilities and the efficiency of laboratory-based evolutionary work.

## MATERIALS AND METHODS

### Strains, plasmids and transformation

The *B. alcalophilus* used in this study was purchased from China General Microbiological Culture Collection Center (CGMCC), while *E. coli* MG1655, haploid *S. cerevisiae* Y187, and diploid *S. cerevisiae* INVSC1 were purchased from Angyu Biotechnology Co., Ltd. (Shanghai, China). The strains and plasmids are listed in Table S1, and the primers are listed in Table S2. Primers, restriction endonucleases, PCR enzymes, and other molecular reagents used for PCR and sequence analysis are sourced from Sangon Biotech (Shanghai,China) Co., Ltd. Firstly, the *BaRecJ* gene (WP_003324372.1) was amplified from the genomic DNA of *B. alcalophilus* through polymerase chain reaction (PCR). Clone the PCR product into pMD18-T using TA cloning method and confirm it through DNA sequencing. Further subcloning *BaRecJ* into *E. coli* expression vectors pGEX-4T-1 (BamHI and XhoI), constructing a full-length BaRecJ expression vector pGEX-BaRecJ, and further constructing different domain expression vectors pGEX-BaRecJ-exo and pGEX-BaRecJ-endo. Based on the codon preference of *S. cerevisiae*, the BaRecJ encoding gene was optimized and nuclear localization signals (NLS) were added at the 5’ and 3’ ends. The gene was cloned into the vector pESC-URA (SalI and HindIII) to construct the *S. cerevisiae* expression vector pESC-BaRecJ. Using heat shock method to transform pGEX-4T-1, pGEX-BaRecJ, pGEX-BaRecJ-exo, and pGEX-BaRecJ-endo into *E. coli* MG1655; Convert pESC-URA and pESC-BaRecJ into *S. cerevisiae* Y187 and INVSC1, respectively, using electroconversion method.

### Media and culture conditions

*B. alcalophilus*, *E. coli*, and *S. cerevisiae* were cultured in BPM (5 g/L peptone, 3 g/L beef extract, 5 g/L NaCl, pH 7.5) medium, LB medium (10 g/L NaCl, 10 g/L peptone, 5 g/L yeast extract), and YPD medium (10 g/L yeast extract, 20 g/L peptone, 20 g/L glucose), respectively. *B. alcalophilus* and *S. cerevisiae* were cultured at 30 L, unless otherwise specified, *E. coli* was cultured at 37 L. The transformants of *E. coli* and *S. cerevisiae* were screened on LB agar plates containing 60 μg/ml ampicillin and SD-Ura agar plates, respectively. In the *E. coli* mutation experiment, 0.2mMIPTG was added as an inducer, and in the yeast mutagenesis experiment, the expression of BaRecJ was induced using SC-Ura medium (with 10 g/L raffinose and 40 g/L galactose added). The fermentation experiment used the fermentation medium reported by Mingming Zhang et al (2017), which contains 4 g/L yeast extract, 3 g/L peptone, and 100 g/L glucose. The batch fermentation was carried out in a 250 ml triangular flask containing 50 ml of fermentation broth under limited aeration conditions Except for the fermentation experiment, all liquid cultures in this study were grown in a 20 mL system at 180 rpm under shaking conditions.

### *E. coli* mutagenesis experiment and Sanger sequencing of *rpoB* gene

Select *E. coli* transformant cells pGEX-4T-1, pGEX-BaRecJ, pGEX-BaRecJ-exo, and pGEX-BaRecJ-endo, and cultivate them until mid-log phase (OD_600_=0.5∼0.8). Subsequently, add IPTG to a final concentration of 0.2mM to the above cultures, and induce cultivation at 16 L to accumulate nucleases under low-temperature conditions. Sample every 4 hours and transfer to 30 L for cultivation, allowing the nucleases to perform cleavage and generate mutations until growth reaches a stationary phase Dilute the optical density of each mutation population mentioned above to 0.3, then perform a 10-fold serial dilution. Plate the diluted bacterial cultures onto LB agar plates and incubate for 24-36 h. Count the CFUs and calculate the Mortality rate of each treatment group. Mortality rate = 1 - (CFUs of the treated group plate / CFUs of the pGEX-4T-1).

Additionally, induce transformed *E. coli* cells in logarithmic phase at 16 L. After sampling every 4 h, transfer the samples to 30 L for plateau phase cultivation. Subsequently, dilute these bacterial samples and plate them onto culture plates containing 100μg/ml rifampicin agar. After incubating for 24-36 h, count the colonies and calculate the relative survival rate of each treatment group. After incubation for 24-36 hours, count the CFUs and calculate the mutation rate for each treatment group, which is represented by the relative survival number. The relative survival number is calculated as the CFUs of the treatment group plate divided by the CFUs of the pGEX-4T-1.

Rifampicin-resistant colonies were selected and inoculated into Eppendorf tubes, followed by overnight growth in LB medium supplemented with 100 µg/mL rifampicin. Overnight culture samples (50 µL) were taken and heated at 100 L for 15 min. PCR amplification was performed with primers AB1 (5′-ATGTCAAATCCGTGGCGTGAC-3′) and AB2 (5′-TTCACCCGGATACATCTCGTCTTC-3′) to amplify *rpoB* fragments. Each fragment was sequenced twice with primers AB3 (5′-CGGAAGGCACCGTAAAAGACAT-3′) and AB4 (5′-CGTGTAGAGCGTGCGGTGAAA-3′).

### Evolution with BaRecJ mutagenesis method

Inoculate the parent strain into YPD media containing varying concentrations of acetic acid (0, 0.1%, 0.2%, 0.3%, 0.4%, 0.5%, 0.6%, 0.7%) and ethanol (2%, 3%, 4%, 5%, 6%, 7%), respectively. Cultivate for 3 days and record the OD_600_ to determine the background tolerance values of the strain to both acetic acid and ethanol.

#### Acetic acid tolerance

Culture the yeast strains Y187-pESC-URA, Y187-BaRecJ, INVSc1-pESC-URA, and INVSc1-BaRecJ, each transformed with different plasmids, until they reach the logarithmic phase. Use SC-Ura medium for centrifugation and resuspension twice to remove glucose. Incubate these cultures in SC-Ura induction medium at 16°C for 5 days to allow for adequate protein accumulation. Transfer the cultures to 30°C to induce mutations, adding inhibitors (0.2% acetic acid for Y187 and 0.4% for INVSc1) and continue cultivation for 3 days. Centrifuge the evolved mutant populations and resuspend them in fresh YPD medium, adjusting the suspension’s OD_600_ to 0.6. Then, inoculate 600 μl of this culture into fresh YPD medium containing (0.3% to 0.8%) acetic acid, incubate for 3 days and record the OD_600_ values. Spread the cultures on YPD agar plates containing ethanol to isolate single clones.

#### Ethanol tolerance

The ethanol evolution experiment follows the same procedure as the acetic acid evolution process. Y187 and INVSc1 use 3% and 6% ethanol, respectively, as the initial stress conditions for the evolutionary populations. Upon completion of the evolution, the cultures are transferred to YPD medium containing ethanol (ranging from 2% to 12%) and then incubated for 3 days to assess cell OD. Subsequently, the cultures are spread on YPD agar plates containing ethanol for the isolation of individual clones

### Fermentation experiments

Inoculate Ia-M2, Ia-08 and INVSc1-BaRecJ (control) into YPD medium, respectively, and culture them with shaking for 12 h, as the seed cultures for fermentation experiments. Inoculate 5 mL of seed culture into a flask containing 45 mL of fermentation medium for batch fermentation. Add 6% ethanol and 0.4% acetic acid to the respective tolerant and control strains. Fermentation conditions are controlled at 30 L and 150 rpm, with natural pH maintained. The residual sugar in the fermentation medium was measured by DNS method to monitor the fermentation process. The residual sugar in the fermentation medium was measured every 24 h, and the fermentation was considered to be finished when the residual sugar content was stable. After the fermentation was finished, the fermentation broth was centrifuged (8000g, 10 minutes), and the supernatant was filtered with a 0.22 µm microporous filter. The analysis was performed by gas chromatography-mass spectrometry instrument (GC-MS) (Agilent, USA) equipped with a DB-624 (60m×250um×1.40um) column, operating at 50 L. High purity helium was used as the carrier gas, with a flow rate of 1mL/min. Each experiment was repeated three times.

### Re-sequencing of whole genome

Ia-M2, Ia-08 and the parental strain INVSc1-BaRecJ were inoculated separately into YPD medium and shaken for 12 h. Then the cells were harvested by centrifugation and the supernatant was removed. For DNA extraction, a MasterPureTM DNA Purification Kit (Epicenter, San Diego, United States) was used, according to the manufacturer’s instructions. The DNA samples were sent to Xi’an Haorui Gene Technology Co., Ltd. (Haorui Genomics, Xi’an, China) for genome resequencing, using PacBio Revio, with an average of 100-fold coverage. For DNA sample quality, the purity of DNA (A260/A280 ratio) was measured using Nanodrop, and precise quantification of DNA was performed using Qubit, ensuring a total DNA amount of ≥ 10µg for each library. DNA samples that passed the quality check were randomly fragmented into 350 bp pieces using the Covaris sonicator. Libraries were constructed using the TruSeq Library Construction Kit. After library construction, initial quantification was conducted using Qubit 2.0, followed by diluting the library to 1 ng/µl. Then, the insert size of the library was assessed using the Agilent 2100. After the insert size met the expected criteria, the effective concentration of the library was accurately quantified using the QPCR method. Raw data were filtered using fastp, and the filtered sequences were aligned to the reference genome SC288C using the BWA alignment software. Trimmomatic v.0.32 was utilized for trimming low-quality bases and residual adapters (Bolger et al., 2014). GeMoMa was used to transfer the annotation of coding sequences from the reference to the *S. cerevisiae* parental strain. The detection of SNPs and Indels variations was carried out using GATK software, and the annotation of these variations was conducted using ANNOVAR software. Raw data were deposited in the NCBI Sequence Read Archive (SRA) database under BioProject PRJNA1060366.

### Data analysis

Prism version 9.0 and SPSS version 19.0 are used for data analysis. Specific statistical tests are noted for individual experiments. All of the GC–MS data had been conveyed by the diluted factor.

## Results

To determine the mutagenic efficiency and application scope of BaRecJ, experiments were conducted in both prokaryotic organisms (*E. coli*) and eukaryotic organisms (*S. cerevisiae*). We constructed expression vectors for *E. coli* MG1655, which contain the full-length sequence of the *BaRecJ* gene (pGEX-BaRecJ), the exonuclease activity domain (pGEX-BaRecJ-exo), and the endonuclease activity domain (pGEX-BaRecJ-endo). We utilized rifampicin resistance as a phenotype to test the Mortality and mutation rates (equal relative survival number) of the full-length and various fragment. (Mortality = 1 - colony forming units (CFUs) in treatment group / CFUs in pGEX-4T-1; relative survival number = CFUs in treatment group / CFUs in pGEX-4T-1). Figure 1A & B display the mutagenic Mortality and mutation rates of each treatment group under different induction times. Both the pGEX-BaRecJ and pGEX-BaRecJ-endo groups exhibited higher Mortality rates, while the pGEX-BaRecJ-exo group showing extremely low cleavage Mortality. The results suggest that the higher mortality rate is primarily due to the endonuclease activity of BaRecJ (Figures 1A). Furthermore, the pGEX-BaRecJ group showed a significantly higher relative mutation rate after 24 h of induction, reaching 199 times that of the empty vector control. In contrast, groups possessing only endonuclease or exonuclease activities exhibited extremely low mutation rates, at 1 and 12 times the empty vector control, respectively. Hence, we can conclude that the efficient mutagenesis by BaRecJ necessitates the concurrent action of both endonuclease and exonuclease activities.

**Figure 1.**
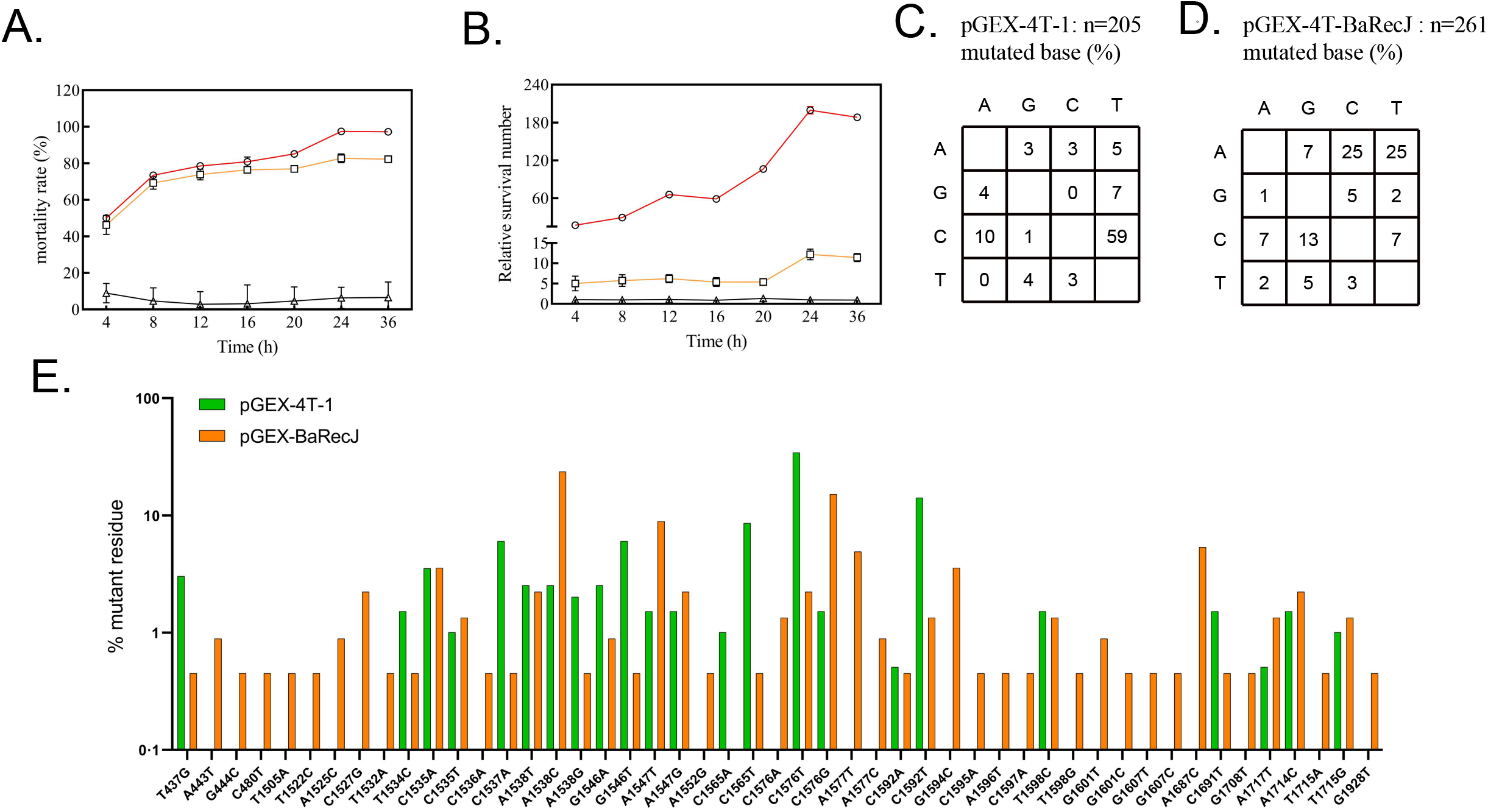
Splicing test of BaRecJ enzyme and mutation spectrum analysis of BaRecJ mutagenesis method. (A-B)The shear mortality rate of different enzyme activity fragments in *BaRecJ* and the relative survival rate on rifampicin (100μg/ml) agar plates. Hollow circle: pGEX-BaRecJ. Hollow triangle: pGEX-BaRecJ-exo. Hollow square: pGEX-BaRecJ-endo. Data are mean±SD (n=3). (C-E) The mutagenic spectra with pGEX-BaRecJ and pGEX-4T-1 (control). Single rifampin-resistant colonies were amplified by PCR and sequenced in *rpoB* gene.

Mutations of the *rpoB* gene are known to endow *E. coli* with resistance to high levels of rifampicin (Badran and Liu, 2015). We selected a rifampicin-resistant mutant population and sequenced the *rpoB* gene to analyze the mutational spectrum. control group (pGEX-4T-1) exhibited a narrow mutagenic spectrum that was strongly biased towards C-T transitions (Figure 1C). However, the experimental group (pGEX-BaRecJ) yielded a broader spectrum with a more uniform distribution of transitions and transversions and a marked increase in the number of A-C, A-T, and C-G transitions than the control (Figure 1D). BaRecJ mutation method obtained 49 different types of base substitution mutations (Figure 1E), significantly higher than mutagenesis methods such as 5-azacytidine (5AZ), ethyl methanesulfonate (EMS), and 2-aminopurine (2AP) (Fernandez et al., 2020).

We constructed a vector suitable for *S. cerevisiae*, in which the nuclear localization signal (NLS) of *S. cerevisiae* was added to upstream and downstream of the *BaRecJ* gene, to target it to the nucleus and enable genome cleavage. We transformed the above vector pESC-BaRecJ and the empty vector pESC-URA into *S. cerevisiae*, and performed mutagenesis experiments. Figures S1 A & B shows the initial acetic acid tolerance of the yeast strains. In the haploid yeast experiment, the acetic acid tolerance phenotype of the control group did not improve, and it could not grow in 0.4% (v/v) acetic acid, while the Y187-BaRecJ was able to adapt to this acetic acid concentration, and its OD_600_ reached 1.3. When the acetic acid concentration increased to 0.6%, the Y187-BaRecJ was still able to grow, and its OD_600_ was 0.64 (Figure 2A). The mutation method based on BaRecJ yield a surprising effect in the adaptation of the diploid INVSc1, the INVSc1-BaRecJ was able to grow in 0.8% acetic acid, while the strains of the control group could not grow even in 0.6% acetic acid (Figure 2B).

**Figure 2.**
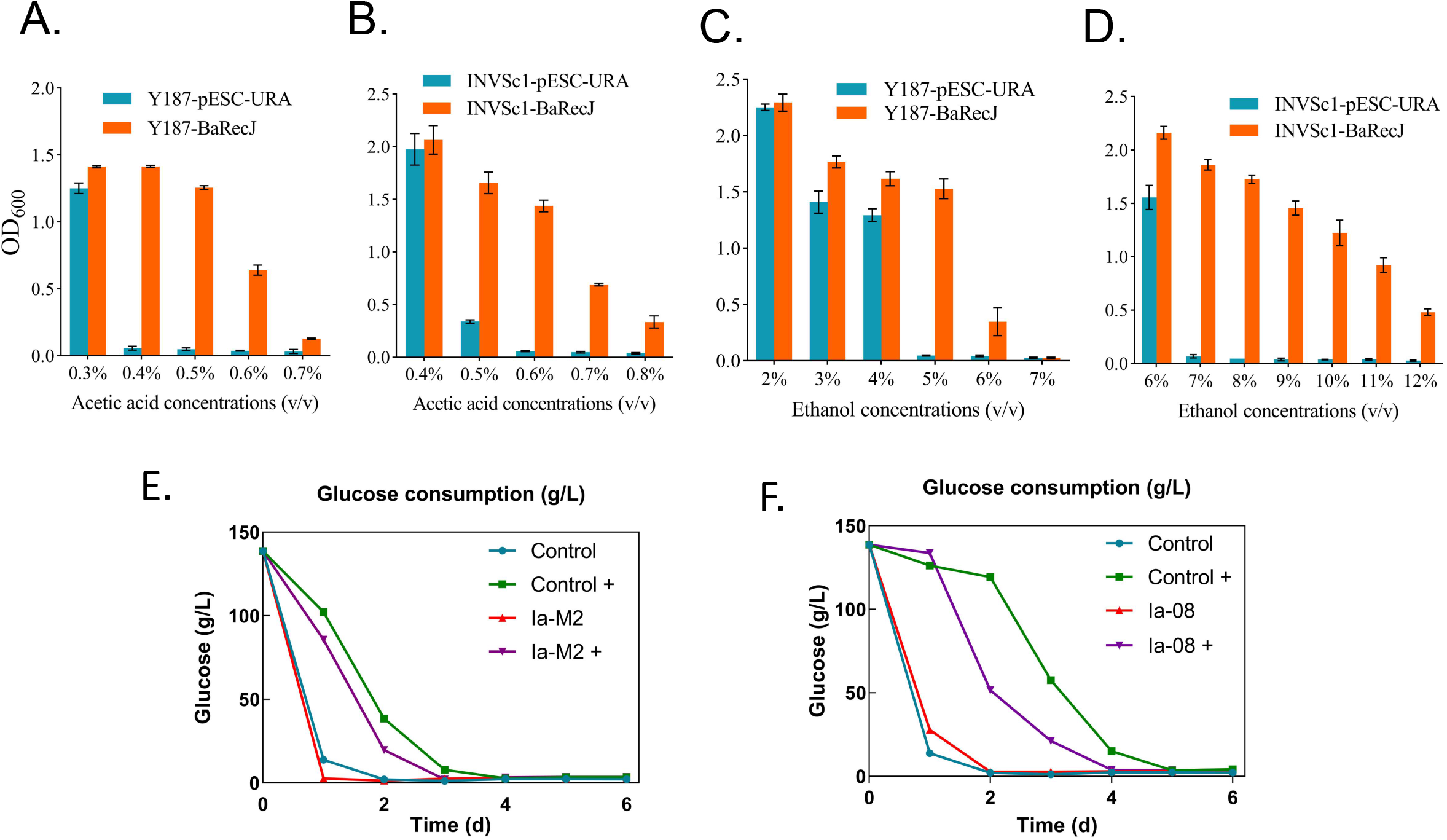
The application of BaRecJ mutagenesis method in improving acetic acid tolerance of Y187 and INVSc1. (A-B) The tolerance of Y187 and INVSc1 mutants to acetic acid. (C-D) The tolerance of Y187 and INVSc1 mutants to ethanol. Data are mean±SD (n=3). Y187-pESC-URA and INVSc1-pESC-URA are Y187 and INVSc1 strains, respectively, transformed with the empty plasmid pESC-Ura. Y187-BaRecJ and INVSc1-BaRecJ are Y187 and INVSc1 strains, respectively, transformed with the vector pESC-BaRecJ. (E-F) Dynamic changes in glucose content during fermentation. (E) Changes in glucose content of mutant strains under 6% ethanol stress and no stress conditions during fermentation. “+” Indicates adding coercive conditions. (F) Changes in glucose content of mutant strains during fermentation under 0.4% acetic acid stress and no stress conditions. “+” Indicates adding coercive conditions. Use INVSC1-pESC-URA as a control. Use DNS colorimetric method to detect residual sugars. Data are mean±SD (n=3).

In the ethanol adaptation experiments of haploid and diploid yeast strains, we also measured the ethanol tolerance background of the yeast (Figures S1 C & D). In the haploid yeast experiment, after of adaptation, the OD_600_ value of the Y187-BaRecJ reached 1.45 at 4% (v/v) ethanol concentration, while the control group was 1.23 (Figure 2C). However, when the ethanol concentration increased to 6%, the growth of the control group was completely inhibited, while the Y187-BaRecJ were still able to grow, and their OD_600_ was 0.35 (Figure 2C). In the diploid experiment, the mutation strategy based on BaRecJ showed a significant effect, the control group could hardly grow in 7% ethanol, and its OD_600_ was only 0.07, while the INVSc1-BaRecJ was able to grow in 7% to 12% ethanol concentrations, and the OD values were 1.87 (7%) and 0.48 (12%), respectively (Figure 2D).

From the populations evolved in ethanol and acetic acid, we isolated two mutants with enhanced tolerance, Ia-M2 and Ia-08, respectively. To evaluate the fermentation performance of the mutant strains, the glucose consumption and fermentation product content of the mutant strains were measured with the parental strain (INVSc1-BaRecJ) as a control. Compared with the control, the glucose consumption ability of Ia-M2 and Ia-08 under non-stress conditions was similar, and the residual sugar content in the medium tended to be stable after three days of fermentation (Figure 2 E & F). Under 6% ethanol stress, the mutant strain Ia-M2 could consume fermentable sugars faster, and almost depleted all the glucose after three days of fermentation, while the control still had 7.8 g/L of residual glucose (Figure 2E). The strain Ia-08 consumed all the glucose after four days of fermentation under 0.4% acetic acid stress, while the control strain still had 15.0 g/L of residual glucose (Figure 2F). The measurement of the main fermentation products showed that, compared with the control, the ethanol production of the ethanol-tolerant mutant Ia-M2 increased by 50.46%, the acetic acid production of the acetic acid-tolerant mutant Ia-08 increased by 71.45%, and the ethanol production also increased by 39.75% (Table. 1) In summary, using the BaRecJ mutagenesis method to domesticate wine yeast can improve the tolerance to acetic acid and ethanol, and improve the fermentation performance.

**Table 1.**

|  | Control | Ia-M2 | Ia-08 |
| --- | --- | --- | --- |
| acetaldehyde (ug/ml) | 83.59 ± 5.63 | 91.65 ± 8.82** | 28.46 ± 2.26** |
| ethanol ( mg/ml ) | 24.84 ± 2.23 | 37.37 ± 0.80** | 34.71 ± 1.14** |
| ethyl acetate (ug/ml) | 5.52 ± 0.48 | 5.94 ± 0.61 | 5.57 ± 0.09 |
| Isobutanol (ug/ml) | 4.03 ± 0.02 | 4.77 ± 0.20 | 3.38 ± 0.44 |
| 3-methyl-1-butanol (ug/ml) | 126.01 ± 8.12 | 116.36 ± 1.55** | 103.91 ± 6.40** |
| acetic acid ( mg/ml ) | 0.66 ± 0.07 | 0.70 ± 0.03 | 1.13 ± 0.02* |
| Isoamyl acetate (ug/ml) | 0.31 ± 0.03 | 0.36 ± 0.02 | 0.58 ± 0.04 |
| benzaldehyde (ug/ml) | 0.23 ± 0.00 | 0.11 ± 0.00** | 0.06 ± 0.00** |
| phenylacetaldehyde (ug/ml) | 0.41 ± 0.05 | 0.35 ± 0.01 | 0.31 ± 0.00 |

Batch fermentation was carried out under non strict anaerobic conditions, and the fermentation supernatant was tested after 5 days. Repeat each experiment three times. One-way ANOVA and Dunnett's post-hoc multiple comparison test were used to evaluate whether the fermentation products of the mutants were different from the parental control. ( Symbols: \**P* value < 0.05; \*\**P* value < 0.01)

Using whole-genome resequencing technology, we compared the genomes of the mutant strains Ia-M2, Ia-08 and the parental strain INVSc1-BaRecJ, and revealed their genomic changes. Based on the reference genome sequence and annotation of *S. cerevisiae* SC288C, we aligned the genomes of the mutant strains and the parental strain. We focused on the detection of the exon regions of the mutant strains, and identified 75 SNPs and 39 Indels [15 insertions (INS), 24 deletions (DEL)] in the exon region of Ia-M2, and 70 SNPs and 39 Indels [16 insertions (INS), 23 deletions (DEL)] in the exon region of Ia-08. According to the reference genome annotation and published information (www.ncbi.nlm.nih.gov), all the mutated genes and the genes related to inhibitor resistance are shown in Table. S3 and Table. 2, respectively..

**Table 2.**
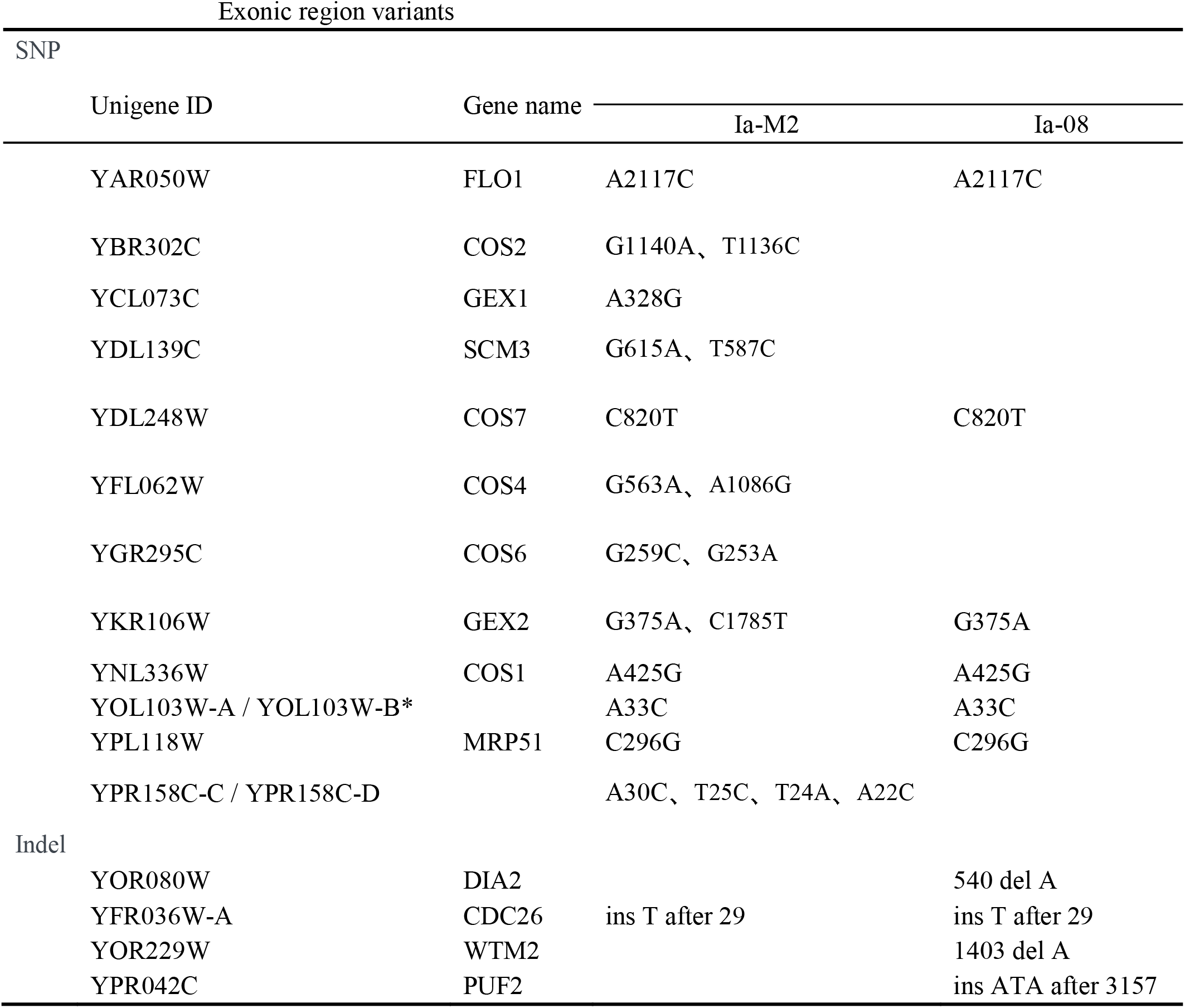
Exonic region variants.

| Exonic region variants |  |  |  |  |
| --- | --- | --- | --- | --- |
| SNP |  |  |  |  |
| Unigene ID | Gene name |  |  |  |
|  |  | Ia-M2 |  | Ia-08 |
| YAR050W | FLO1 | A2117C |  | A2117C |
| YBR302C | COS2 | G1140A、 T1136C |  |  |
| YCL073C | GEX1 | A328G |  |  |
| YDL139C | SCM3 | G615A、 T587C |  |  |
| YDL248W | COS7 | C820T |  | C820T |
| YFL062W | COS4 | G563A、 A1086G |  |  |
| YGR295C | COS6 | G259C、 G253A |  |  |
| YKR106W | GEX2 | G375A、 C1785T |  | G375A |
| YNL336W | COS1 | A425G |  | A425G |
| YOL103W-A / YOL103W-B* |  | A33C |  | A33C |
| YPL118W | MRP51 | C296G |  | C296G |
| YPR158C-C / YPR158C-D |  | A30C、 T25C、 T24A、 A22C |  |  |
| Indel |  |  |  |  |
| YOR080W | DIA2 |  |  | 540 del A |
| YFR036W-A | CDC26 | ins T after 29 |  | ins T after 29 |
| YOR229W | WTM2 |  |  | 1403 del A |
| YPR042C | PUF2 |  |  | ins ATA after 3157 |

## Discussion

Here, we present a novel global in *vivo* mutagenesis approach, characterized by a broader mutation spectrum, capable of generating a more genetically diverse mutant population, thereby yielding mutants with superior traits. This is attributed to the dual genomic cleavage processes of BaRecJ. One process may involve endonuclease-induced genomic breaks, triggering an SOS response, followed by repair via non-homologous end joining. The other process initiates with endonuclease-induced genomic breaks, followed by exonuclease-mediated degradation starting from the ends, leading to mutations during the polymerase-mediated repair phase. The BaRecJ mutagenesis method is a unique mutagenesis mechanism that differs from the current global in *vivo* mutagenesis methods. It is not necessary to have a deep understanding of the mismatch repair genes of the target strain, and to construct plasmids with defective DNA polymerase, simply transfer BaRecJ into recipient cells for in vivo mutagenesis (Xu et al., 2018; Luan et al., 2013).

The sequencing of the rifampicin resistance gene *rpoB* in *E. coli* showed that this mutagenesis method produces the most mutation sites among all global mutagenesis methods. Based on this method, we performed a mutagenesis on *E. coli* and obtained 49 types of mutations, while the MP6 mutagenesis method is the one that produces the most types of mutations among other methods, only 32 (Badran and Liu, 2015). In addition, other mutagenesis methods obtain fewer mutation sites, including 5-azacytidine (5AZ) (10), ethyl methanesulfonate (EMS) (10), 2 aminopurine (2AP) (14), cisplatin (CPT) (8) and ultraviolet (UV) light (10), yielded only 8 - 14 types of base substitution mutations (Fernandez et al., 2020). This further demonstrates that the BaRecJ mutagenesis method can produce a broader spectrum, and has greater potential in global mutagenesis.

In the Ia-M2 and Ia-08 mutants, we detected mutations in GEX1 and GEX2, which have solute: proton antiporter activity and can mediate H^+^ efflux. Their mutations may accelerate H^+^ efflux, enabling the mutants to tolerate higher concentrations of ethanol or acetic acid (Zhang et al., 2017). *S. cerevisiae* can use cell surface mannose groups to connect with adjacent cells and form flocs to cope with stress, and the FLO family plays an important role in this process (Hodgson et al., 1985; Katju et al., 2009; Rossouw et al., 2015 Ye et al., 2022). We detected a mutation in FLO1 in Ia-M2 and observed significant flocculation(Figure S2), which provides help in studying how *S. cerevisiae* can improve tolerance based on flocculation (Ye et al., 2022).

Ethanol and acetic acid are crucial factors affecting the fermentation process in brewing yeast, sharing similar inhibitory mechanisms. Upon entering the cell, they induce acidification in both the cytoplasm and vacuolar environment (Ma and Liu., 2010; Arneborg et al., 2000). This acidification impacts protein structures, leading to misfolding and intracellular aggregation, causing cellular damage and functional disruptions (Wang et al., 2023). In mutants Ia-M2 and Ia-08, mutations were detected in genes such as COS1, COS2, COS4, COS6, COS7, SCM3, DIA2, and CDC26. These genes participate in ubiquitin-dependent protein degradation, and their mutations might accelerate the degradation of denatured proteins induced by acidification, reducing cellular harm. Additionally, changes were observed in genes associated with protein transcription and translation. Mutations were detected in certain transcription and translation-related genes in the mutants, such as YOL103W-A, YOL103W-B, MRP51, YPR158C-C, YPR158C-D, WTM2, and PUF2. These variations in genes might aid in promoting the synthesis of new proteins. We speculate that Ia-M2 and Ia-08 likely renew intracellular proteins by degrading denatured proteins and synthesizing new ones to sustain their activity, countering the impact of cellular acidification on normal metabolism, it is a new mechanism for tolerating acetic acid and ethanol.

In summary, we have developed a novel and original biotechnological technique for random mutagenesis, with a mutagenic mechanism entirely distinct from any other current random mutagenesis methods. This technology offers a broader spectrum and higher efficiency and can be extensively applied to the breeding of higher organisms. It is of significant importance for enriching mutant libraries, studying gene functions, and conducting metabolic engineering for breeding purposes.

## Supporting information

raw data of rpob

## Acknowledgement

This work was supported by funding from the National Key Research and Development Program of China (2018YFC0310701).

Minggang Zheng, Jixaing Shang, and Yanchao Zhang conducted experiments and performed data analyses; Experimental materials provided by Zhongtao Sun and Minggang Zheng; Minggang Zheng, Jixaing Shang, Zongjun Xu and Shouqing Zhang prepared the manuscript. All authors approved the final version of the manuscript.

---

**Fig S1.**
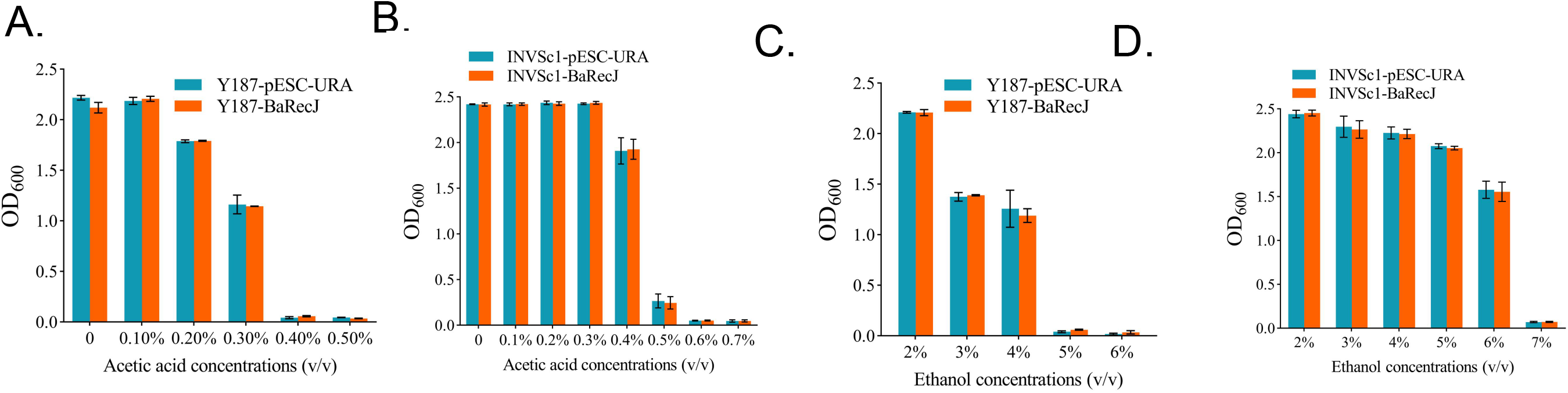
Tolerance background of Y187 and INVSC1. (A-B) Background of acetic acid tolerance in 187-pESC-URA, Y187-BaRecJ, INVSC1-pESC-URA, INVSC1-BaRecJ. (C-D) Background of ethanol tolerance in 187-pESC-URA, Y187-BaRecJ, INVSC1-pESC-URA, INVSC1-BaRecJ. Data are mean±SD (n=3). Y187-pESC-URA and INVSc1-pESC-URA are Y187 and INVSc1 strains, respectively, transformed with the empty plasmid pESC-Ura. Y187-BaRecJ and INVSc1-BaRecJ are Y187 and INVSc1 strains, respectively, transformed with the vector pESC-BaRecJ.

**Figure S2.**
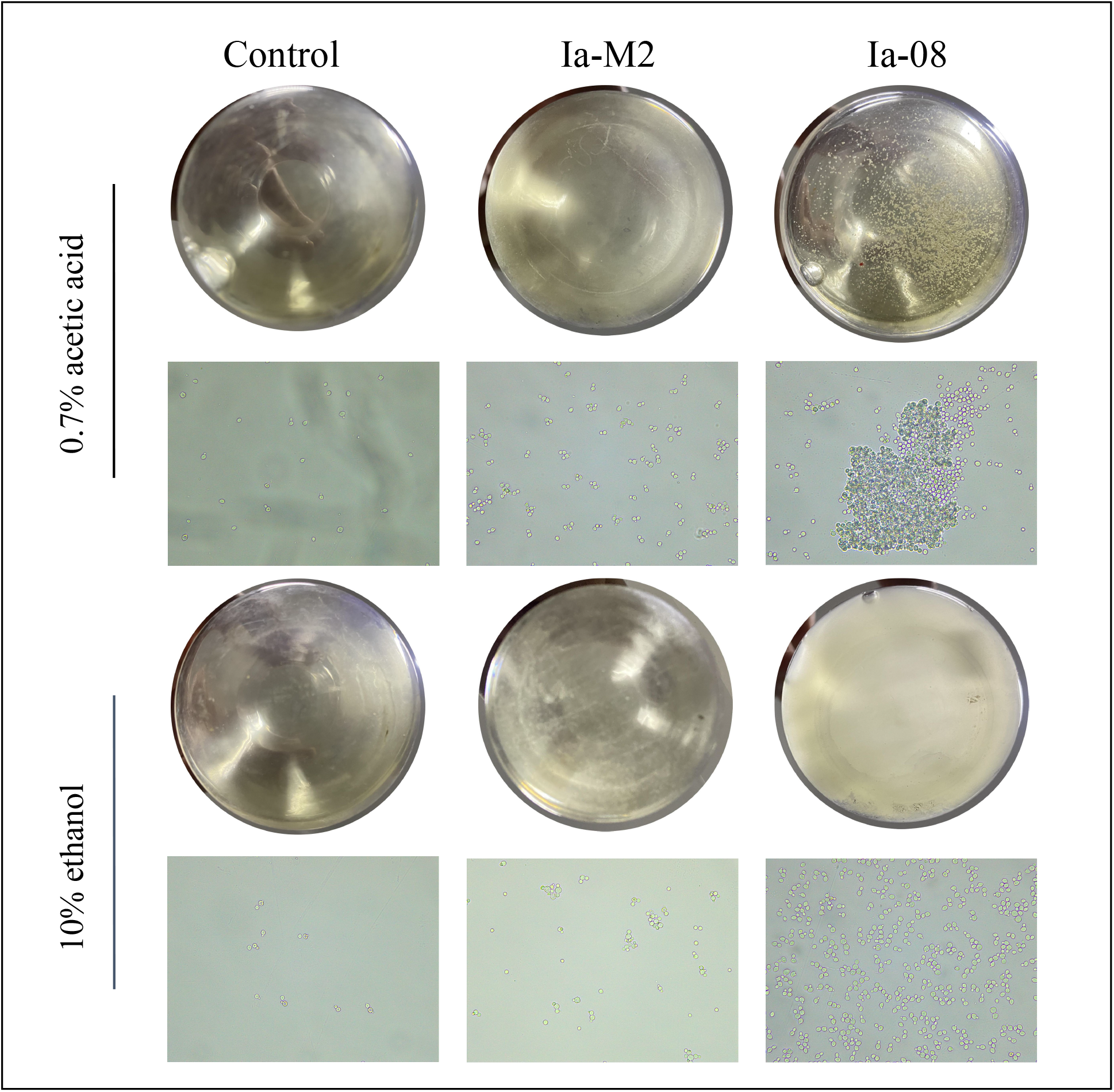
Phenotypic observation of mutant strains. Observation of flocculating strain and non-flocculating strain INVSc1-BaRecJ. Upper panel, photograph of the shake flask culture; lower panel, images under optical microscope.Using the parent strain as a control, culture for 2 days in media containing 0.7% acetic acid and 10% ethanol, respectively. Observing cell phenotype with naked eye and optical microscope (magnification 10× 40).

