## Supplementary material for "Highly efficient in *vivo* mutagenesis method based on the endonuclease and exonuclease activity of *Bacillus alcalophilus* RecJ (BaRecJ)": Table S1 Table S2 & TableS3: Table S1.docx

Strains and plasmids

| **Strains,** **plasmids**  **and** **primers** | **Relevant characteristics** | **Source** |
| --- | --- | --- |
| **Strains** | | |
| *Bacillus. alcalophilus* | Purchased from the China General Microbiological Culture Collection Center (CGMCC) | CGMCC 1.3604 (=ATCC27647) |
| E. *coli* MG1655 | K12 F- λ- ilvG- rfb-50 rph-1 | AngYu Bio |
| *S. Cerevisiae* Y187 | MATα, ura3-52, his3-200, ade 2-101, trp 1-901, leu 2-3,  112, gal4Δ, met–, gal80Δ, URA3 : : GAL1UAS-GAL1TATAlacZ, MEL1 |  |
| *S. Cerevisiae* INVSc1 | MATa his3Δ1 leu2 trp1-289 ura3-52/MATα his3Δ1 leu2  trp1-289 ura3-52 |  |
| **Plasmids** | | |
| pMD18-T | ColE1 origin, Ap^R^ | TaKaRa Bio |
| pGEX-4T-1 | pMB1 origin, Ap^R^ |  |
| pGEX-BaRecJ | pMB1 origin, Ap^R^ |  |
| pGEX-BaRecJ-  exo | pMB1 origin, Ap^R^ |  |
| pGEX-BaRecJ-  endo | pMB1 origin, Ap^R^ |  |
| pESC-Ura | pUC origin, Ap^R^, 2-micron origin, Ura3, 6.6kb | AngYu Bio |
| pESC-BaRecJ | pUC origin, Ap^R^, 2-micron origin, Ura3 |  |

**Abbreviations:** Cm, chloramphenicol; Ap, ampicillin; Kan, Kanamycin; R, resistance
