## Supplementary material for "Highly efficient in *vivo* mutagenesis method based on the endonuclease and exonuclease activity of *Bacillus alcalophilus* RecJ (BaRecJ)": Table S1 Table S2 & TableS3: Table S2.docx

| Plasmid | Oligos | Sequence(5’-3’) | Template | Backbone | Restriction sites |
| --- | --- | --- | --- | --- | --- |
| pBaRecJ | BaRecJ-f1 | ATAAACCGTGGAGCGAGACC | Genomic DNA of *Bacillusalc alophilus* (ATCC 27647 and CGMCC 1.3604) | pMD18-T | / |
|  | BaRecJ-r1 | CCACCAGTTGCCAATAAGTC |  |  |  |
|  | BaRecJ-f2 | ATGTTGAATCCAAAGGCAAG |  |  |  |
|  | BaRecJ-r2 | TTATACCGTCTCCTTGATTG |  |  |  |
| pGEX-BaRecJ | pGEX-BaRecJ-f | CACACAGGAAACAGTATTCATGTTGAATCCAAAGGCAAG | pBaRecJ | pGEX-4T-1 | BamHI/XhoI |
|  | pGEX-BaRecJ-r | CCCGGGAATTCCGGGGATCCTTATACCGTCTCCTTGATTG |  |  |  |
| pGEX-BaRecJ-exo | pGEX-BaRecJ-exo-r | CCCGGGAATTCCGGGGATCCTTATTGTTTAATAATATCTAAAG |  |  |  |
| pGEX-BaRecJ-endo | pGEX-BaRecJ-endo-f | CACACAGGAAACAGTATTCATGAAACAACTTGCACCAT |  |  |  |
| pESC-BaRecJ | pESC-BaRecJ-r | TAATACGACTCACTATAGGGCCCGGGCGTCGACATGGATTATAAAGATGACGATG | Gene Synthesis | pESC-Ura | SalI/HindIII |
|  | pESC-BaRecJ-f | AGCGGATCTTAGCTAGCCGCGGTACCAAGCTTTTAGACCTTTCTCTTCTTTTTTGGAGGT |  |  |  |

Primers
